# Sexual tail dimorphism explains speciation rate in swallows and martins

**DOI:** 10.1101/2024.12.11.627946

**Authors:** Masaru Hasegawa

## Abstract

Sexual selection can promote speciation in theory, but macroevolutionary studies have reported mixed results. A possible explanation for this inconsistency is the use of inappropriate proxies of sexual selection. Here, by focusing on swallows and allies (Aves: Hirundinidae), I examined whether or not a classic example of sexually selected trait, long outermost tail feathers, explains speciation rate in this clade. Long tails have been repeatedly shown to be intersexually selected in manipulative experiments, which is further corroborated by a series of macroevolutionary studies, thereby validating their use as a target of sexual selection. I found that hirundines with sexually dimorphic tail length have a significantly higher speciation rate than those with sexually monomorphic tail length. Furthermore, evolutionary changes in the extent of sexual tail dimorphism (and those in the extent of male outermost tail exaggeration) was significantly positively associated with speciation rate. Sexual plumage dichromatism and wing dimorphism are irrelevant to speciation rate. Together, the current study demonstrated the importance of using valid targets of sexual selection in studies on the macroevolutionary patterns of speciation. Using a classic, unidimensional sexual trait (i.e., long tail), I demonstrated a strong macroevolutionary support that divergence of intersexually selected traits promotes speciation.

## Introduction

Why some, but not all, clades exhibit lineage diversification is a major issue in evolutionary biology (Wien 2017). Since the influential work of Darwin (e.g., Darwin 1871), sexual selection is thought to facilitate speciation and thus lineage diversification as well, because intersexually selected traits and their diversification can promote reproductive isolation and increase reinforcement at secondary contact (e.g., West-Eberhard 1983; Panhuis et al. 2001; Ritchie 2007 for some reviews). However, macroevolutionary studies have reported mixed results, with some studies supporting this theory (e.g., Ellis & Oakley 2016; Portik et al. 2019; Price-Waldman et al. 2020) whereas others not supporting or reporting more-nuanced results (e.g., Huang & Rabosky 2014; Cooney et al. 2017; Miller et al. 2021). Overall, the patterns supporting theory are often restricted to certain clades (e.g., certain bird family), and macroevolutionary analyses using wider phylogenies (e.g., all Aves) have often failed to support the theory (e.g., Kraaijeveld et al. 2011; Huang & Rabosky 2014; Miller et al. 2021).

An explanation for the observed inconsistency (and the absence of global support throughout the tree of life) is the difficulty of choosing appropriate proxies of sexual selection. For example, sexual dimorphism is a major proxy, but its validity is often questioned for at least three reasons (e.g., Janicke et al. 2018; Cally et al. 2021; Miller et al. 2021). First, besides sexual selection, which is predicted to promote speciation (see the preceding paragraph), several other processes, including intersexual niche segregation and sex differences in resource availability, can drive the evolution of sexual dimorphism. Thus, the mere presence of sexual dimorphism might be a poor predictor of sexual selection (and, conversely, sexual monomorphism in conspicuous traits does not preclude the presence of intense sexual selection on traits inconspicuous to human observers: Cally et al. 2021). Not surprisingly, the importance of intersexual selection (and intrasexual selection) cannot be elucidated from these studies. Second, proxies of sexual selection (e.g., sexual plumage dichromatism) often contain multiple components (e.g., melanin-based, carotenoid-based, depigmented, structural coloration, and their combination in separable body regions such as the head, body, and flight feathers), each of which have its own ecological functions (e.g., antioxidant, parasite resistance, and structural strength: see Hill & McGraw 2006a,b for bird coloration). Hence, although sexual selection is sometimes considered a “non-ecological” driver of speciation, the benefits and costs of sexual traits depend on ecology (e.g., Maan & Seehausen 2011; Mendelson & Safran 2021). Accordingly, even if the focal proxy of sexual selection (e.g., sexual plumage dichromatism) evolved through sexual selection, the ecology of species affects the expression of each component (and hence total expression as well), again rendering the proxy a poor correlate of sexual selection across species. For example, sexual plumage dichromatism in carotenoid-based coloration would be affected not only by sexual selection but also by the availability of carotenoids, which cannot be synthesize in birds (Hill & McGraw 2006a,b; also see McGraw 2008 for similar ecological constraints on melanin-based coloration). As a consequence, the association between the proxy of sexual selection and speciation should be weakened when the focal taxa (e.g., Aves) exhibit a variety of ecological background. Finally, most previous studies have tested the association between the proxies of sexual selection and speciation assuming that intense sexual selection promotes speciation without paying attention to the target traits, but, rather, divergent sexual traits should promote speciation (e.g., Gomes et al. 2016; Price-Waldman et al. 2020). This means that researchers should focus on the phenotypic changes of the “targets” of sexual selection in the focal lineage. In this context, using appropriate proxies of targets of sexual selection is crucial (e.g., Price-Waldman et al. 2020). Ideally, qualitatively homologous, single-component sexually selected traits should be chosen as a proxy among species sharing similar ecological background (e.g., foraging niche, locomotion, and mating system), so that we can avoid blurring a relationship between sexual selection and speciation due to the confounding effects of multiple sources of selection and ecological background (which affects each component of sexual traits in different manners; see preceding sentences in this paragraph). Easily measurable, unidimensional traits are particularly well-suited, because they can avoid the difficulty of quantifying complex traits accompanying multiple axes (e.g., how visible and ultraviolet hue, saturation, brightness, and color pattern should be integrated in complex, integument color signals). However, it is often difficult to find out such clades.

The family Hirundinidae (or hirundines, including swallows and allies) is a suitable study system for investigating speciation in relation to sexual selection. Hirundinidae includes a model species for studying sexual selection, i.e., the barn swallow *Hirundo rustica*, in which sexually dimorphic, long outermost tail feathers are repeatedly shown to be the target of intersexual selection (e.g., Møller 1988; reviewed in Møller 1994; Romano et al. 2017; Hasegawa 2018) and a driver of reproductive isolation between subspecies (Schield et al. 2024). Tail length can be easily evaluated along a single axis across species (i.e., short–long axis) and thus can avoid the difficulty of quantifying complex traits accompanying multiple axes (see the preceding paragraph). Although long outermost tail feathers that protrude from the short central tail feathers, thus making “forked tails” were once considered an “aerodynamic device” for efficient aerial foraging (e.g., Evans 1998), the manipulative experiment used to validate this hypothesis was logically flawed and thus inconclusive (see Hasegawa 2024 for details). Although the function of slightly longer outermost tail feathers than central tail feathers (i.e., shallowly forked tails) remains unclear (e.g., Hasegawa 2024), a series of recent macroevolutionary studies consistently support that long outermost tail feathers have a foraging cost (e.g., Hasegawa et al. 2016; Hasegawa & Arai 2017a, 2018a,b, 2022) and coevolved with proxies of the intensity of sexual selection, including opportunities for extra-pair mating (e.g., Hasegawa & Arai 2020a, 2022; Hasegawa 2023). At the least, the aerodynamic device hypothesis has no explanation for sexual dimorphism in tail length, and hence sexually dimorphic tail length can be used as a proxy of sexual selection, as it evolved with opportunities for extra-pair mating (Hasegawa & Arai 2020a). Furthermore, hirundines share similar ecology, including hyper-aerial insectivores, typically socially monogamous with biparental provisioning (Turner & Rose 1994), which is suitable for studying the effect of sexual selection on speciation without being confounded by a variety of ecological background (see the preceding paragraph).

In the current study, I examined speciation rate in relation to sexual dimorphism in tail length in hirundines (family: Hirundinidae) using macroevolutionary analyses, specifically, the State-dependent Speciation and Extinction framework (i.e., SSE; Maddison et al. 2007). The presence of sexual selection would promote speciation, and hence I expected that the presence of sexual dimorphism in tail length would be positively associated with speciation rate. To study whether the observed patterns are unique to the proxy of sexual selection used here (i.e., sexual dimorphism in tail length), I also tested alternative proxies of sexual selection, namely, sexual dimorphism in wing length as a measure of sexual dimorphism in body size (see Hasegawa & Arai 2018b; also see Methods for details), and sexual plumage dichromatism, i.e., a well-known proxy of sexual selection (see above). In addition, to examine how sexual selection affects speciation, I focused on the extent of sexual dimorphism in tail length and its evolutionary changes. Recent macroevolutionary studies of birds demonstrated that evolutionary changes in ornamentation, rather than the extent of ornamentation (used as a proxy of intensity of sexual selection), affects speciation (e.g., Gomes et al. 2016; Price-Waldman et al. 2020), and thus I expected that this would also be the case in the current study. Although the importance of trait change is not unique to sexual traits (as similar patterns can be expected in non-sexual traits, including species recognition traits; Wien 2017; Price-Waldman et al. 2020), confirming the combination of the two predictions, i.e., 1) positive association between the presence of intense sexual selection (or, in practice, its proxy) and speciation, and 2) positive association between the changing expression of sexual traits and speciation rate, would provide better support for the importance of sexual selection in the context of speciation.

## Methods

### Data collection

As before (e.g., Hasegawa & Arai 2017a, 2018b, 2020a,b), information on sexual dimorphism in wing length and sexual dimorphism in tail length was obtained from Turner & Rose (1994). I used the monograph (Turner & Rose 1994), because this publication provides detailed information on hirundines compared with books dealing with wider taxa (e.g., all birds). When subspecies differences were found, the description of the nominate subspecies was used. Sexual dimorphism in wing length was used as a proxy for sexual dimorphism in body size, because wing length is tightly correlated with other measurements of body size (e.g., body mass) and does not impose a reduction of sample size (see detailed information in Hasegawa & Arai 2018b, 2020a). Sexual dimorphism in tail length used in the current study (i.e., sex difference in tail length: e.g., Hasegawa & Arai 2020a) roughly matches a >5% sex difference, which is the reference value generally considered to classify a species as sexually monomorphic/dimorphic (e.g., Cuervo & Møller 1998, 1999; Aparcio et al. 2003; also see Table S1 for the dataset). An exception is *Hirundo neoxena*, which is described to have sexually dimorphic tails (but no sexually dimorphic measurements). However, when *H. neoxena* is considered to have sexually dimorphic tails, the results of SSE did not change (i.e., significant and nonsignificant results remained unchanged; data not shown), and hence this inconsistency exerted at best a minor effect on the results (also see the Results section for evolutionary changes of male outermost tail length proportional to central tail length in relation to speciation rate, which does not depend on sexual monomorphism/dimorphism classification). The same index of sexual tail dimorphism was found to be coevolved with opportunities for extra-pair mating (Hasegawa & Arai 2020a), supporting the use of this binary variable as a proxy of sexual selection (i.e., this variable reflects sexual selection at least in part). In addition, I calculated male tail length relative to female tail length as the extent of sexual tail dimorphism to estimate a potential predictor of speciation, evolutionary change of ornamentation (see the Statistics section for details).

I also recorded any male–female plumage color discrepancy, i.e., a measure of sexual plumage dichromatism, which was collected from the “Descriptions” section in the literature (Turner & Rose 1994). This criterion was chosen to be comparable with sexual dimorphism in tail length (i.e., >5% sex difference, as explained above) rather than much clearer sexual plumage dichromatism as noted in the plate section (e.g., Hasegawa & Arai 2020a; note that clear sexual plumage dichromatism is relatively rare in hirundines, i.e., 11/72 = 15%, and hence is impractical to yield unbiased estimates, though using this proxy did not change the results qualitatively; details not shown; Table S1). Finally, opportunities for extra-pair mating, measured as incubation type (i.e., female-only vs. biparental incubation), was also obtained mainly from Turner & Rose (1994), as done in previous studies (e.g., Hasegawa & Arai 2020a, 2022). However, I could collect data from 41 species of 72 hirundine species (i.e., 56%), which might affect the accuracy and precision of the estimation of speciation and extinction rates (FitzJohn et al. 2009), and hence the results are not reported here. The data set for the current study is summarized in Table S1 and Fig. S1.

## Statistics

I used BiSSE (i.e., Binary State-dependent Speciation and Extinction) framework to analyze state-dependent speciation and extinction model (Maddison et al. 2007). For this purpose, I used the R package “diversitree” (Fitzjohn et al. 2009), and I used sexual dimorphism in tail length as the binary state variable (see above). Because the BiSSE model might show “false positive” under the presence of rate shifts independent of the traits of interest (e.g., Rabosky & Goldberg 2015), I also used the package “hisse” (i.e., the Hidden State-dependent Speciation and Extinction; Beaulieu & O’Meara 2016), which can account for the rate shifts by including “hidden states” (i.e., to detect rate shifts independent of the traits of interest, and, if required, to account for them). I thus first compared models with and without hidden states and confirmed that character-independent models (i.e., CID models) with hidden states demonstrated poor fit to the data, justifying the use of the BiSSE framework (i.e., no apparent rate shifts independent of the traits of interest; see Results). An alternative, nonparametric method, FiSSE (i.e., acronym of the Fast, Intuitive, State-dependent Speciation and Extinction analysis; Rabosky & Goldberg 2017), was not used in the current study due to the modest sample size (i.e., 72 species), as FiSSE has limited statistical power for phylogeny with fewer than 300 species. Thus, as explained in the next paragraph, I instead used ES-sim analysis, which is an alternative approach to study trait-associated speciation rate (reviewed in Morlon et al. 2024). A total of 1000 alternative trees of Hirundinidae were downloaded from birdtree.org (Jetz et al. 2012; also see Sheldon et al. 2005 for original phylogeny on swallows), which was sufficient to control for phylogenetic uncertainty (Rubolini et al. 2015). I first compared HiSSE models, in which models with ΔAIC >2, i.e., difference in Akaike information criterion (AIC) values between the best model and the focal model, indicate no substantial support (Burnham & Anderson 2002). Consequently, I confirmed substantial support for the BiSSE model over models with hidden states. Thus, to obtain accurate estimation, I estimated speciation rate, extinction rate, and net diversification rate (i.e., speciation rate − extinction rate) in relation to sexual tail dimorphism using the supported model with all the 1000 trees. The “mcmc” function of “diversitree” was used to obtain 1000 samples for each tree, and then all mcmc (i.e., acronym of Markov-Chain Monte Carlo) outputs of models from 1000 trees were summed up to examine the association between sexual selection and speciation. Following Revell & Harmon (2017), mcmc sampling was initiated with exponential prior and maximum likelihood estimates for each tree for rapid convergence. Because of skewed posterior distribution (see Results), I used 95% credible intervals (CIs) based on highest posterior density, and, when the CIs did not include zero, the estimate was regarded as “significant” (e.g., when the 95% CI of estimated difference in speciation rates between sexually monomorphic and dimorphic species did not include zero, the difference was regarded as significant; see Results). Each statistical test was conducted based on an *a priori* prediction (i.e., sexual tail dimorphism as a proxy of sexual selection was predicted to affect speciation rate), and thus it is not necessary to correct for multiple testing (e.g., see Perneger 1998; Matsunaga 2007; Rubin 2021 for the rationale). In addition, I used sexual dimorphism in wing length and sexual plumage dichromatism as alternative proxies of sexual selection (see Introduction) and repeated the analyses. In the analysis of sexual plumage dichromatism, however, the maximum likelihood estimates for some trees did not converge, and thus mcmc sampling was initiated instead with the maximum likelihood estimates of one randomly chosen tree (tree#1, here).

Finally, to further verify the relationship between sexual tail dimorphism and speciation rate, I used ES-sim analyses (the R function, “essim”: Harvey & Rabosky 2018). This is a tip-rate correlation test and utilize simulation to test the association between continuous traits and speciation rates, measured as a simple statistics, inverse equal split, derived from the branch lengths of a phylogenetic tree (e.g., see Price-Waldman et al. 2020 for its application in the context of sexual selection and speciation). For this purpose, I focused on the extent of sexual tail dimorphism, measured as log((male tail length)/(female tail length)), i.e., the ratio of male and female tail length (note that no hirundine species has female tail length longer than male tail length; Turner & Rose 1994). The use of this proportional index is justifiable as recent studies have demonstrated that mate choice is often based on proportional difference rather than absolute difference (e.g., Caves & Kelly 2023), although this index is mathematically equivalent to sex differences accounted for interspecific variation in body size, i.e., log(sex difference in tail length divided by female tail length +1) and hence the current study cannot be used to distinguish whether animals use proportional or absolute differences. Based on this index, the evolutionary rates (i.e., evolutionary changes) of the extent of sexual tail dimorphism were estimated (Fig. S2), using the function “RRphylo” in the package “RRphylo” (Castiglione et al. 2018), which can account for the rate heterogeneity of continuous traits among individual lineages rather than entire clades (e.g., see Price-Waldman et al. 2020 for its application in the context of sexual selection and speciation). Using the 1000 phylogenetic trees, I calculated mean estimates as the representative values (i.e., I used equal weight for 1000 phylogenetic trees to obtain global mean estimates across trees). Because the extent of sexual tail dimorphism theoretically depends on male and female tail length, I repeat analyses using male outermost tail feather length in proportion to their central tail feather length, i.e., log((male outermost tail feather length)/(male central tail feather length)), and the corresponding female value, i.e., log((female outermost tail feather length)/(female central tail feather length)), to separate the effect of male and female trait expression. All analyses were conducted in R ver. 4.4.2 (R Core Team 2024).

## Results

Analyses of a series of HiSSE models (i.e., BiSSE, CID-1, CID-2, CID-3, and CID-4) revealed that the BiSSE model with sexual tail dimorphism as the binary state of interest fit substantially better (i.e., ΔAIC > 2) than the CID models with and without hidden states (i.e., CID-1 to CID-4; Table 1) in 990 out of 1000 alternative trees. In the remaining 10 trees, CID-1 was chosen as the best model, in which only two cases showed ΔAIC > 2 (2.59 and 3.98; others: range: 0.05–1.97). It should also be noted that the CID models with hidden states (i.e., CID-2 to CID-4) always fit worse than the CID-1 model having no hidden states (i.e., ΔAIC > 2; Table 1), indicating no apparent rate shift independent of the sexual tail dimorphism in this clade (i.e., no support for “hidden” states). Thus, I concentrated below on the analyses of focal models without hidden states (i.e., BiSSE models; note that CID-1 is the special case of BiSSE where speciation and extinction are assumed to be independent of sexual tail dimorphism).

**Table 1.** Model fit of alternative models for state-dependent speciation and extinction using 1000 alternative trees of hirundines (family: Hirundinidae)

| Model | Mean log Likelihood [range] | k | Mean AIC [range] | Mean ΔAIC [range] |
| --- | --- | --- | --- | --- |
| BiSSE | -251.52 [-278.62, -211.73] | 6 | 511.04 [431.46–565.23] | 0.02 [0.00–3.98] |
| CID-1 | -252.77 [-280.34, -213.41] | 4 | 517.54 [438.82–572.68] | 6.51 [0.00–13.65] |
| CID-2 | -258.37 [-285.17, -218.87] | 5 | 526.73 [447.74–580.34] | 15.70 [8.60–23.12] |
| CID-3 | -258.00 [-284.85, -218.50] | 7 | 529.99 [450.99–583.70] | 18.97 [12.50–29.60] |
| CID-4 | -257.81 [-284.71, -218.33] | 9 | 533.63 [454.66–587.43] | 22.60 [16.27–31.63] |

Detailed information of each model is described below (also see Methods):
BiSSE: Binary state-dependent speciation and extinction without hidden states, with sexual tail monomorphism/dimorphism as a binary state variable.
CID-1: character-independent speciation and extinction model without hidden states
CID-2: character-independent speciation and extinction model with a hidden state
CID-3: character-independent speciation and extinction model with two hidden states
CID-4: character-independent speciation and extinction model with three hidden states
k indicates the number of parameters in each model.
AIC indicates Akaike information criterion, and ΔAIC indicates the difference in AIC values between the best model and focal model

The BiSSE model showed that lineages with sexually dimorphic tails have a significantly higher speciation rate than lineages with sexually monomorphic tails in hirundines (lineages with sexual tail monomorphism: mean [95% CI] = 0.09 [0.03–0.15]; lineages with sexually dimorphic tails: 0.25 [0.13–0.41]; mean difference [95% CI] = 0.16 [0.03–0.34]; Fig. 1). On the other hand, there was no detectable difference in extinction rate between them (lineages with sexually monomorphic tails: 0.03 [0.00– 0.11]; lineages with sexually dimorphic tails: 0.06 [0.00–0.18]; mean difference [95% CI] = 0.03 [−0.10 to 0.18]; Fig. 1). As a consequence, the model showed that lineages with sexually dimorphic tails have a marginally higher net diversification rate (i.e., speciation rate − extinction rate) than lineages with sexually monomorphic tails (mean difference [95% CI] = 0.14 [−0.02 to 0.33]; Fig. 1). Moreover, the model showed that the net diversification rate of lineages with sexually dimorphic tails, but not that of lineages with sexually monomorphic tails, is significantly higher than zero (lineages with sexually monomorphic tails: 0.05 [−0.04 to 0.13]; lineages with sexually monomorphic tails: 0.19 [0.08–0.32]), indicating the effect of sexual tail dimorphism (or its close correlate) on lineage diversification.

**Fig. 1.**
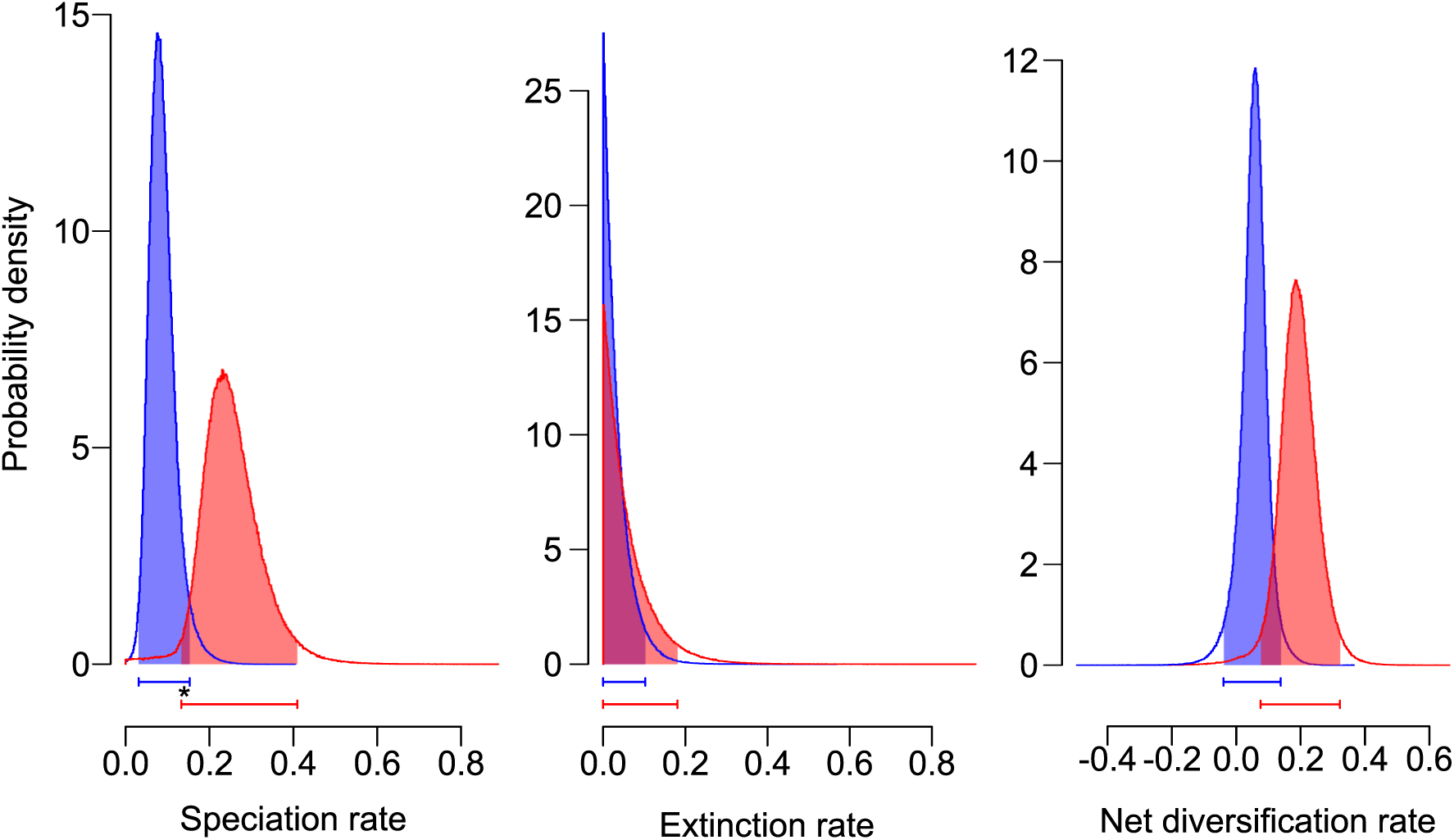
Posterior distribution for speciation rate (left column), extinction rate (middle column), and diversification rate, deduced from speciation rate minus extinction rate (right column) of the species with and without sexual tail dimorphism (depicted by red and blue, respectively) in the swallows and martins (family: Hirundinidae). Bars under each histogram indicate 95% CIs of each estimate and asterisk (*) indicates significant difference between them (see text for formal analyses)

Sexual dimorphism in wing length, a measure of sexual size dimorphism (see Methods), was not significantly associated with speciation rate (mean difference [95% CI] = 0.06 [−0.11 to 0.25]), extinction rate (0.05 [−0.07 to 0.25]), and diversification rate (0.01 [−0.20 to 0.21]; Fig. S3), indicating that the abovementioned finding does not reflect sexual dimorphism in body size. Similarly, sexual plumage dichromatism was not significantly associated with speciation rate (mean difference [95% CI] = 0.02 [−0.22 to 0.29]), extinction rate (0.04 [−0.08 to 0.21]), and diversification rate (−0.03 [−0.32 to 0.29]; Fig. S4), reinforcing the finding that not all types of sexual dimorphism are associated with speciation and extinction patterns. In these analyses, the diversification rate was significantly larger than zero in sexually monomorphic species (lineages with sexually monomorphic wings in the analysis of sexual wing dimorphism: 0.11 [0.03– 0.18]; lineages with sexually monochromatic plumage in the analysis of sexual plumage dichromatism: 0.12 [0.02–0.22]). This was not the case in sexually dimorphic species (lineages with sexually dimorphic wings in the analysis of sexual wing dimorphism: 0.12 [−0.05 to 0.28]; lineages with sexually dichromatic plumage in the analysis of sexual plumage dichromatism: 0.09 [−0.13 to 0.33]; Figs. S3–S4), which is in contrast to the results of sexual tail dimorphism (see above). Concerning sexual plumage dichromatism, qualitatively similar results were found when excluding minor sex differences (Fig. S4; details not shown).

When the relationship between the extent of sexual tail dimorphism and speciation rate was tested using 1000 trees, no significant correlation was observed between them (mean value: rho = 0.14, P = 0.54; Fig. 2 left panel). In contrast, the absolute evolutionary rate (i.e., evolutionary change) of the extent of sexual tail dimorphism was significantly positively correlated with the speciation rate across the 1000 trees (mean values: rho = 0.51, P < 0.01; Fig. 2 right panel). Qualitatively similar patterns were observed between the male outermost tail feather length in proportion to their central tail feather length and speciation, wherein not the extent of male outermost tail exaggeration (mean values: rho = 0.23, P = 0.28) but its absolute evolutionary change was significantly positively correlated with the speciation rate (mean values: rho = 0.42, P = 0.03). Female tail length exhibited similar but weaker relationships (mean values: extent: rho = 0.23, P = 0.29; change: rho = 0.39, P = 0.06), as predicted by intersexual genetic correlation.

**Fig. 2.**
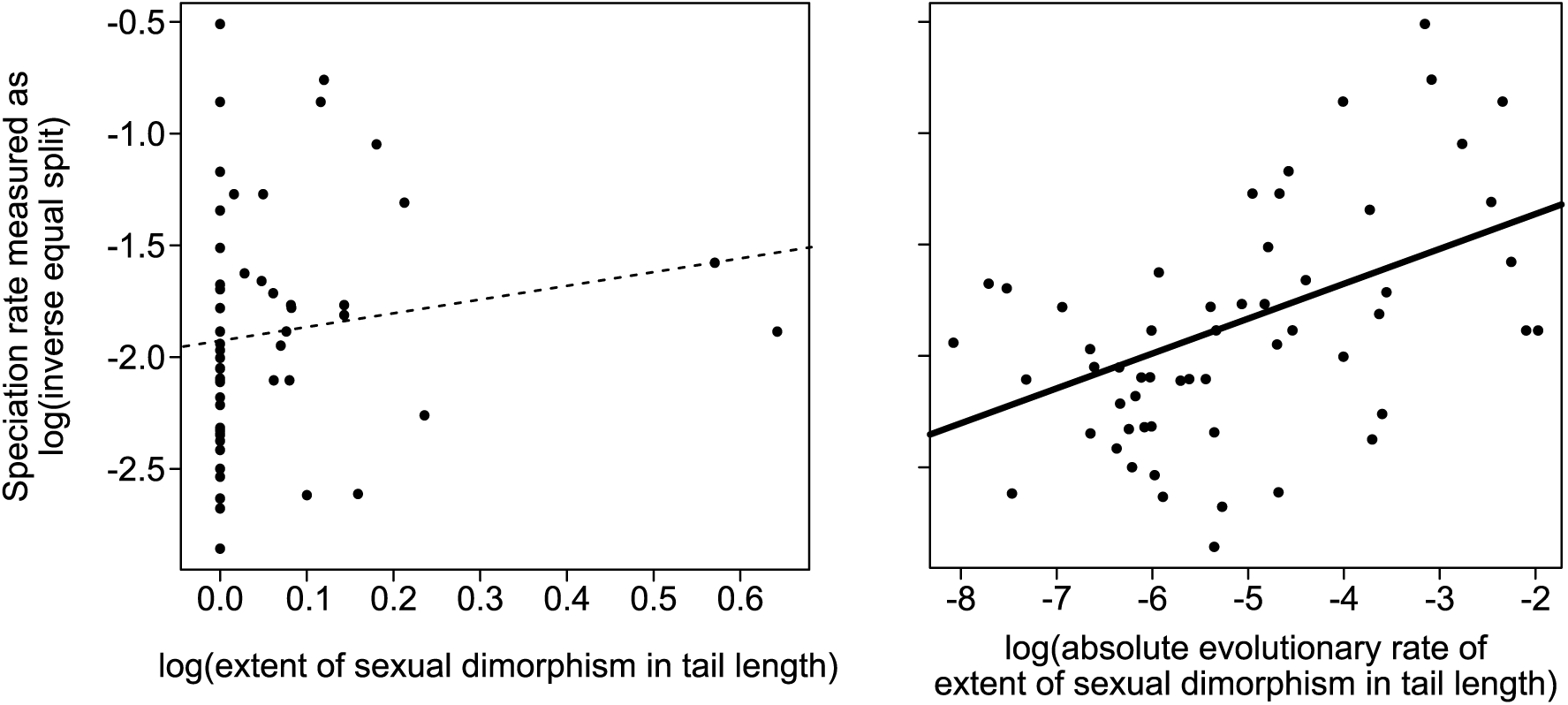
Speciation rate, measured as log(inverse equal split), in relation to the extent of sexual dimorphism in tail length, measured here as log((male tail length)/(female tail length)), as depicted in the left panel and in relation to the evolutionary change of the extent of sexual dimorphism in tail length, estimated by absolute evolutionary rates in log((male tail length)/(female tail length)) using the R function “RRphylo” in the package “RRphylo” after log-transformation (see text), as depicted in the right panel. Here, estimates using one of 1000 phylogenetic trees with simple regression lines were depicted for illustrative purpose (broken and solid lines were used for non-significant and significant correlations; see text for formal analyses using 1000 trees)

When the extent of sexual dimorphism in wing length, instead of the extent of sexual tail dimorphism, was examined in relation to speciation rate, no significant correlation was observed between them (mean values: rho = −0.06, P = 0.80; Fig. S5 left panel). This was also the case when the absolute evolutionary change of the extent of sexual wing dimorphism was examined in relation to speciation rate (mean values: rho = 0.03, P = 0.79; Fig. S5 right panel). Qualitatively similar patterns were observed when male wing length and their evolutionary change were used (mean values: log(male wing length): rho = 0.10, P = 0.60; log(female wing length): rho = 0.13, P = 0.56; absolute evolutionary rate of log(male wing length): rho = 0.32, P = 0.15; absolute evolutionary rate of log(male wing length): rho = 0.34, P = 0.12).

## Discussion

The main finding of the current macroevolutionary study is that hirundines with sexually dimorphic tails have a higher speciation rate than hirundines with sexually monomorphic tails. As the extinction rate was shown to be relatively small with no detectable difference between them, hirundines with sexually dimorphic tails, but not hirundines with sexually monomorphic tails, have a positive diversification rate. These findings are in contrast to those obtained using alternative proxies of sexual selection, namely, sexual dimorphism in wing length and sexual plumage dichromatism, and hence, by comparing the results of alternative proxies, the current study demonstrated the importance of choosing appropriate proxies, or “targets,” of sexual selection, which was further reinforced by the positive association between evolutionary changes in the extent of sexual tail dimorphism and speciation rate. Although trait-independent proxies of sexual selection are sometimes suggested in recent days to test the importance of sexual selection on speciation (e.g., Bateman’s gradient: e.g., Janicke et al. 2018), trait-dependent proxies provide additional information (i.e., how sexually selected traits are associated with speciation, here), which helps us infer the mechanisms underlying the relationship, as I discuss below.

The finding that not the extent of sexual tail dimorphism but its evolutionary change was positively associated with the speciation rate (Fig. 2; also see the Results section for a similar positive link between the change in the extent of male outermost tail exaggeration and speciation rate) is consistent with previous studies using other bird families, wherein evolutionary changes in potential sexual traits are positively associated with the speciation rate (e.g., Gomes et al. 2016; Price-Waldman et al. 2020). This pattern makes sense as the divergence in sexual traits, rather than the absolute expression of sexual traits, would promote speciation. Moreover, unlike most other studies, which use composite traits (e.g., sexual plumage dichromatism across body regions) and thus do not distinguish intersexual and intrasexual selection (and possible other causes of sexual dimorphism; see Introduction), the current study focused on a classic unidimensional intersexually selected trait (with no known function in intrasexual selection; Møller 1994; also see Schield et al. 2024 for its effect on reproductive isolation between subspecies), thus providing a strong support that intersexual selection promotes speciation. Although how evolutionary changes in sexual traits are associated with speciation rate remains to be clarified (e.g., pre-zygotic and post-zygotic reproductive isolation, and reinforcement), divergence in sexual selection was observed even within species in hirundines (e.g., Safran & McGraw 2004; Romano et al. 2017), suggesting that not only post-zygotic isolation through reduced viability of hybrids (e.g., via mismatch between tail length and the compensatory traits; Hasegawa 2024) but also pre-zygotic reproductive isolation (and reinforcement) through differential mating success would be involved, which needs to be clarified in future studies.

A caveat of the current study is that I could not completely exclude potential confounding factors. For example, the current proxy of sexual selection, sexual tail dimorphism, does not preclude the involvement of additional sexual traits, such as song. In fact, a previous study demonstrated the synergistic coevolution of song and forked tails in hirundines (Hasegawa 2023). Thus, changing extent of sexual tail dimorphism might accompany the changes in song, contributing to their association with speciation rate, although this possibility remains to be tested. Likewise, the extent of sexual tail dimorphism might be negatively correlated with the use of other unknown sexual traits (e.g., cryptic sexual traits might be used in short-tailed species), further promoting speciation through divergent sexual selection. In fact, even in single species, e.g., the barn swallow, many types of sexual signals are reported (reviewed in Hasegawa 2018), and hence it is likely that multiple ornaments in combination affect mating and reproductive success (e.g., see Hasegawa & Arai 2017b for a negative interplay of tail length and plumage coloration at pair formation), potentially influencing reproductive isolation. Finally, there exists a possibility that speciation is driven not by sexual traits but by some non-sexual traits that are closely linked to the sexual traits; however, currently, it is not known whether and how evolutionary changes in the extent of sexual tail dimorphism (and male tail ornamentation) track such traits (except for changing benefits and costs of tail length).

In summary, I demonstrated here that hirundines with sexually dimorphic tails have a higher speciation rate than hirundines with sexually monomorphic tails, suggesting the importance of sexual selection in speciation, although the correlational nature of the current study cannot completely exclude the possibility that sexually dimorphic tails are strongly related to an unmeasured driver of speciation. Rather than using sexual overall dimorphism or sexual plumage dichromatism as rough proxies of sexual selection (while ignoring all other factors contributing to sexual dimorphism) to increase apparent sample size across clades (e.g., Aves), focusing on the targets of sexual selection in a given clade would be fruitful to clarify whether and how sexual selection promotes speciation. The targets of sexual selection would depend on the ecology of each clade (e.g., aerial foraging in hirundines, which affects the costs, and possibly benefits, of tail ornamentation), and such ecology-dependent sexual selection can answer the question why the association between certain proxies of sexual selection and speciation is not global but clade-specific, which should not be dismissed in future studies.

## Acknowledgments

I thank Dr Emi Arai for valuable comments and thank Dr Shumpei Kitamura and his lab members at Ishikawa Prefectural University for their kindest advices. I am grateful to Dr Angela Turner for her kindly support on the valuable information on swallows.

## Data availability

The data sets supporting this article have been uploaded as part of electronic supplementary material, table S1.

## Author contribution

I performed the data analysis and wrote the manuscript alone.

## Declarations

### Ethics approval

All information were obtained from literature and no animal subjects were used in the current study.

## Supplementary Materials

**Table S1.**
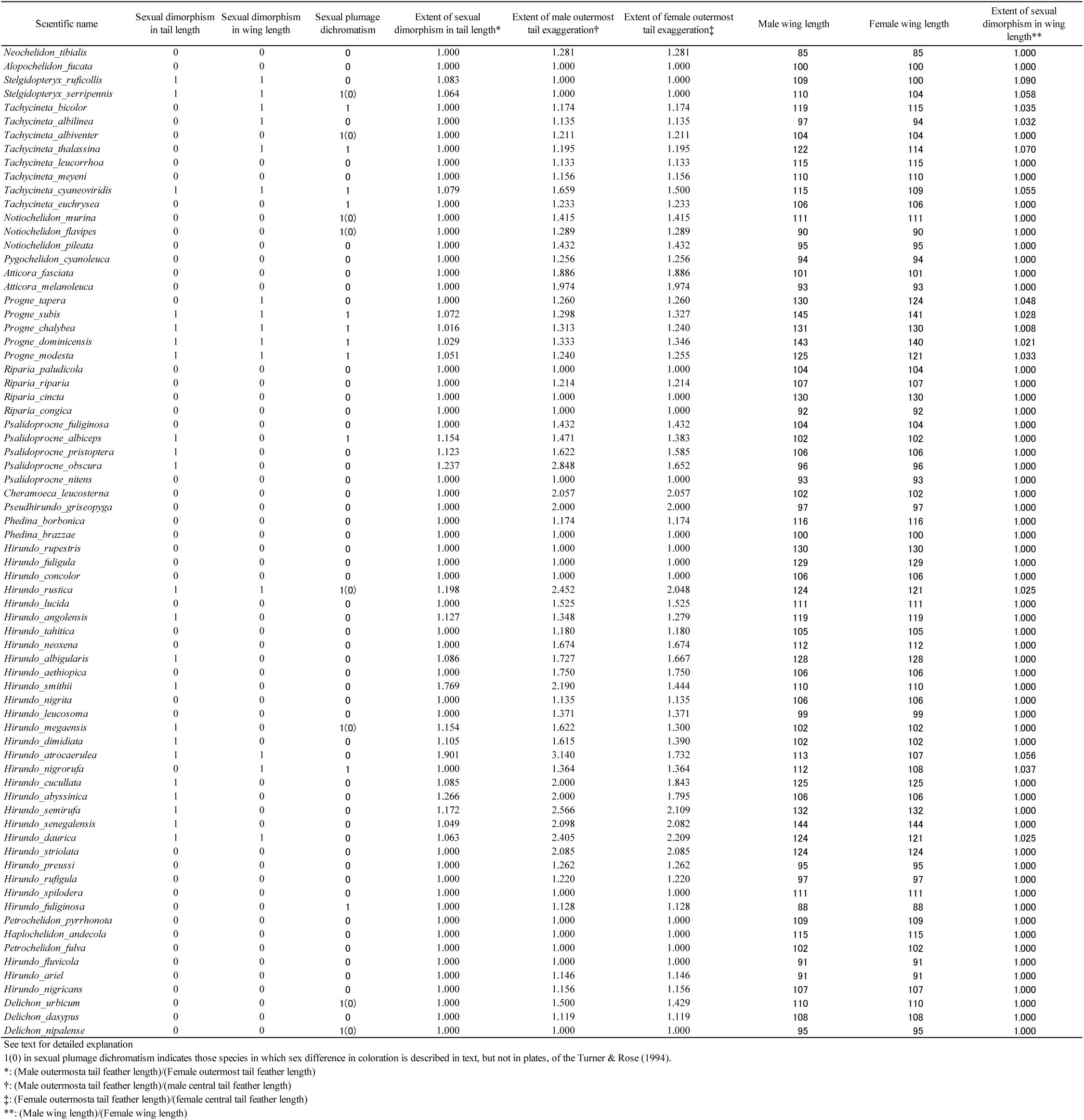
Dataset for the current study (*n* = 72; see text for the detailed methods)

**Fig. S1.**
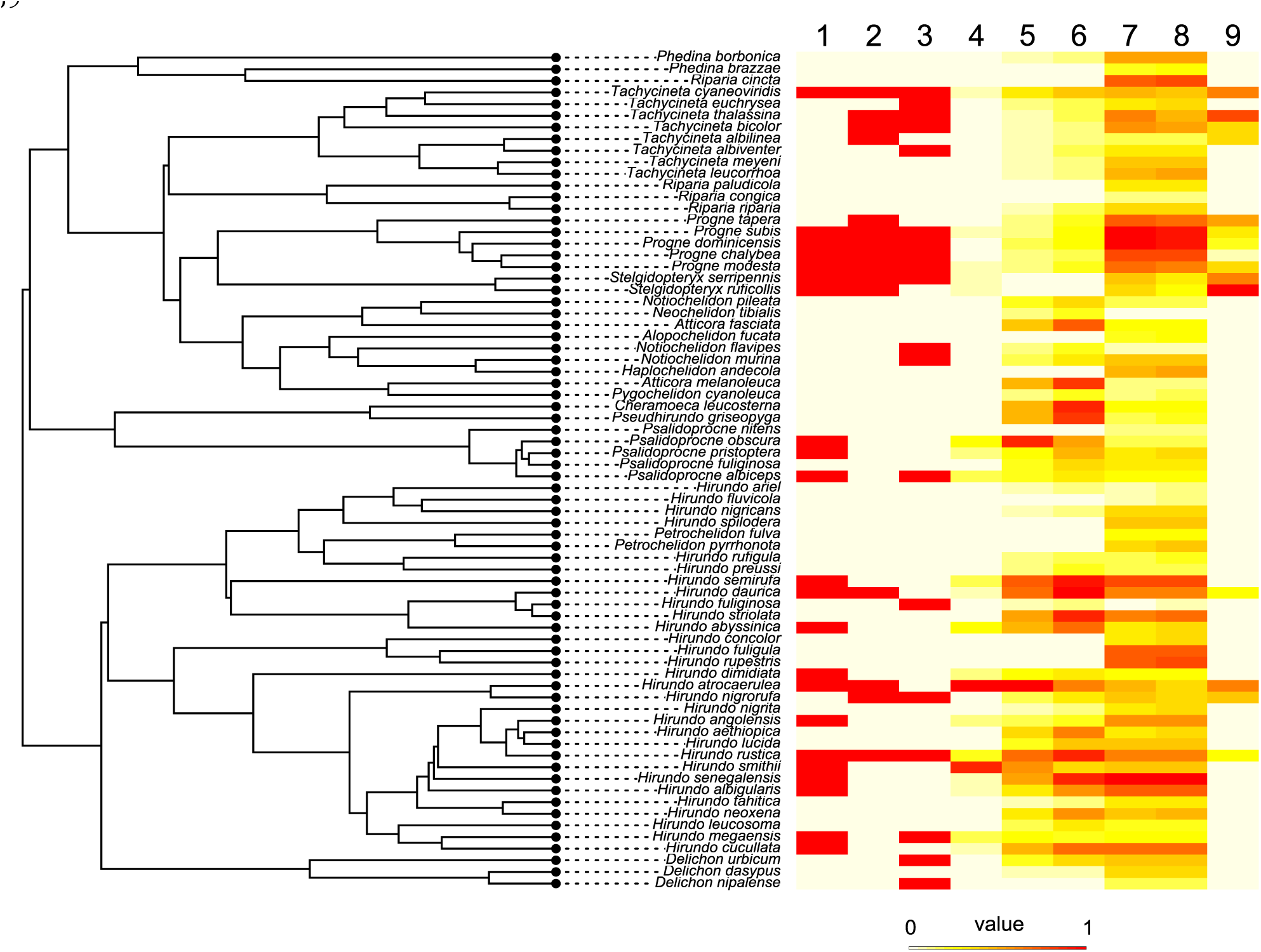
An example of phylogenetic tree of the family Hirundinidae (n = 72) obtained from birdtree.org. Besides the tree, we plotted nine traits, indicated by no. 1‒9, using the function “phylo.heatmap” from the package “phytools” (Revell 2012); 1: sexual dimorphism in tail length; 2: sexual dimorphism in wing length; 3: sexual plumage dichromatism; 4: extent of sexual dimorphism in tail length; 5: extent of male outermost tail exaggeration; 6: extent of female outermost tail exaggeration; 7: male wing length; 8: female wing length; 9: extent of sexual dimorphism in wing length). Each trait was normalized using the min-max normalization to be comparable across traits (see Table S1 for detailed explanation for each trait)

**Fig. S2.**
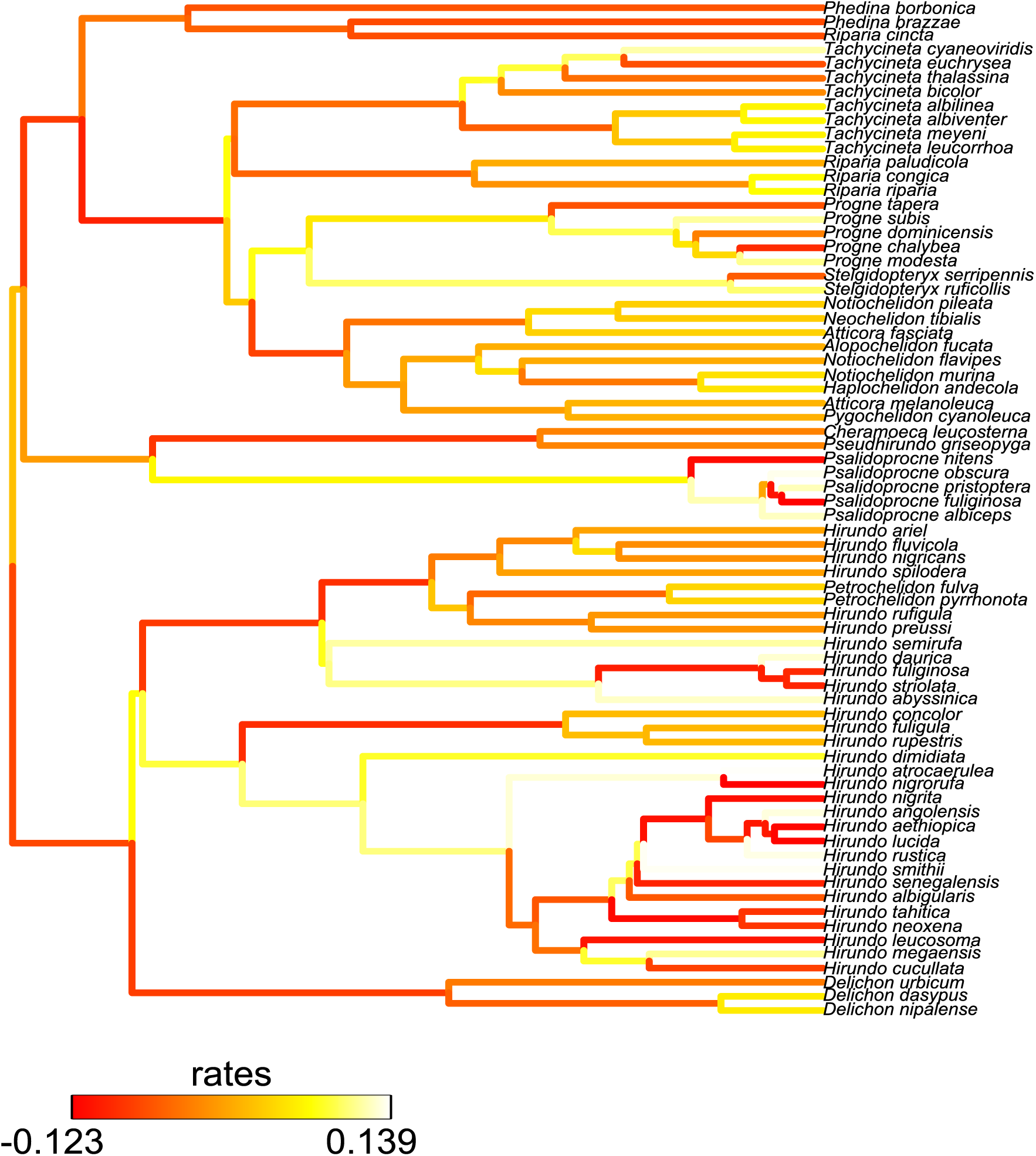
An example of evolutionary rates of the extent of sexual dimorphism in tail length, estimated by using the function “RRphylo” in the package “RRphylo” (see text for detailed information). Because the estimate depends on phylogenetic trees, evolutionary rate of the extent of sexual dimorphism in tail length per each tree was estimated to analyze its relationship with speciation rate (see text)

**Fig. S3.**
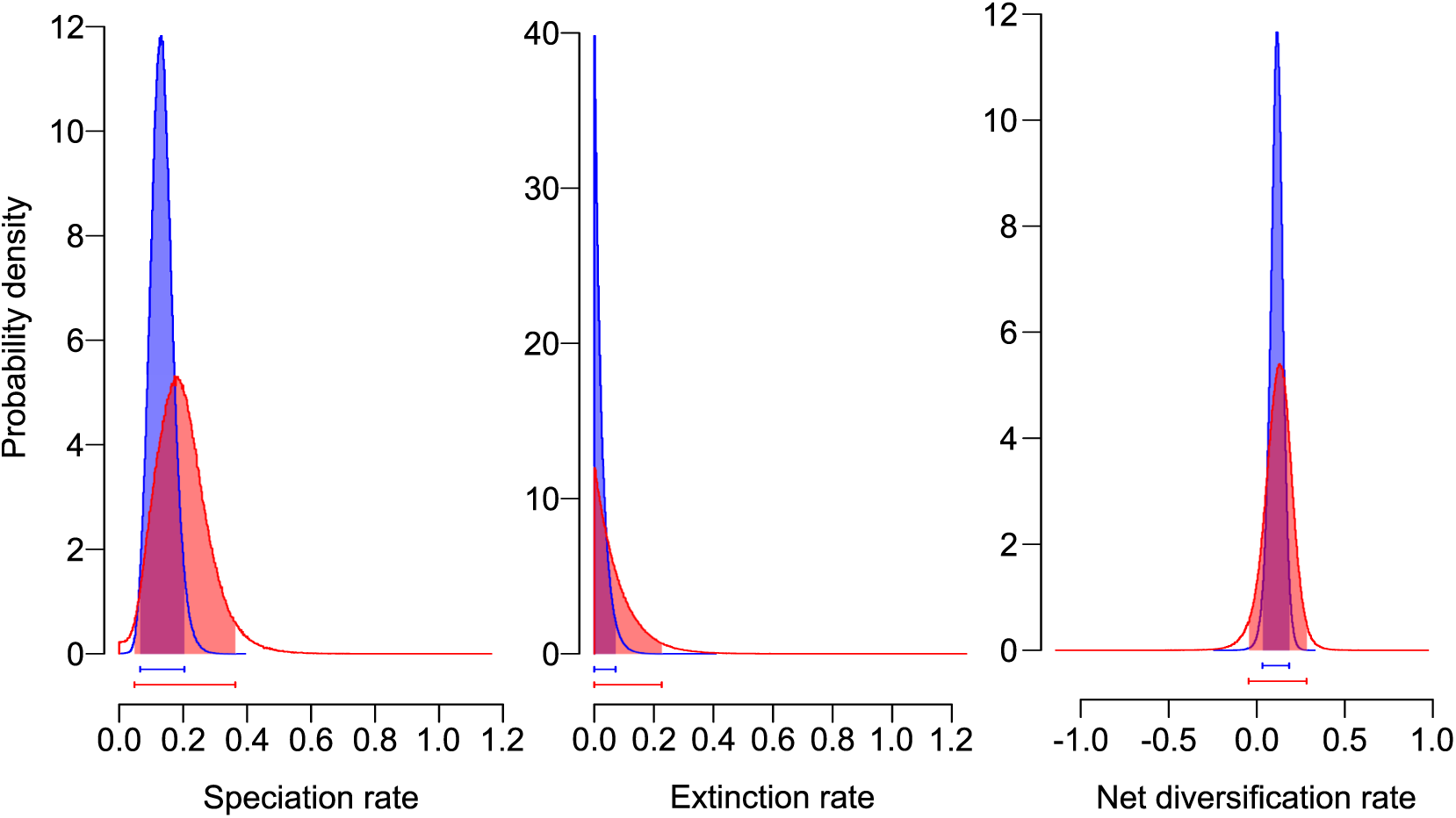
Posterior distribution for speciation rate (left column), extinction rate (middle column), and diversification rate (right column), deduced from speciation rate minus extinction rate of the species with and without sexual wing dimorphism (depicted by red and blue, respectively) in the swallows and martins (family: Hirundinidae). Bars under each histogram indicate 95% CIs of each estimate (see text for formal analyses)

**Fig. S4.**
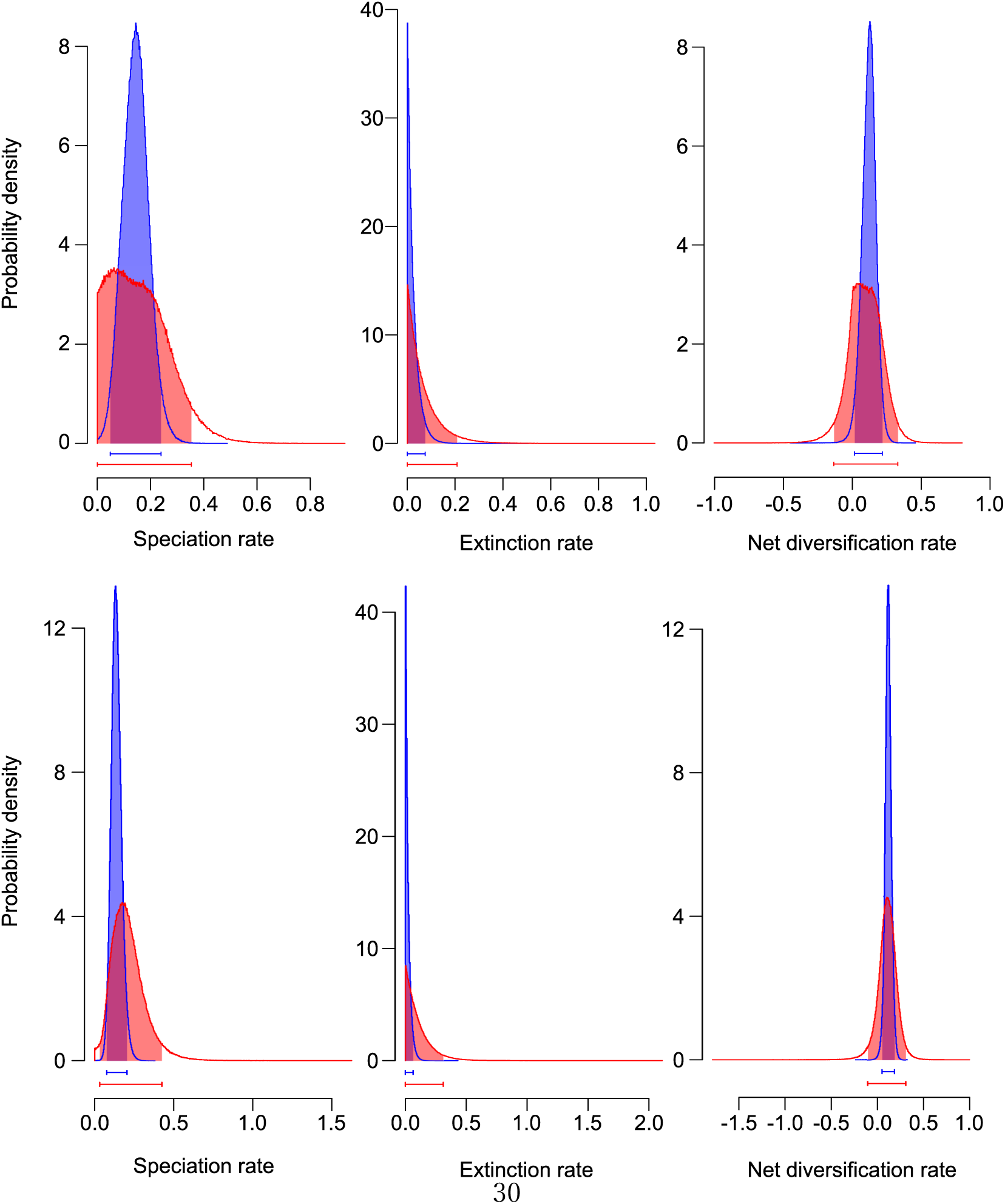
Posterior distribution for speciation rate (left column), extinction rate (middle column), and diversification rate (right column), deduced from speciation rate minus extinction rate of the species with and without sexual plumage dichromatism (depicted by red and blue, respectively) in the swallows and martins (family: Hirundinidae): upper raw: all sex differences in coloration described in Turner & Rose (1994) were included; lower raw: only sex differences described in plates in Turner & Rose (1994) were included. Bars under each histogram indicate 95% CIs of each estimate (see text for formal analyses)

**Fig. S5.**
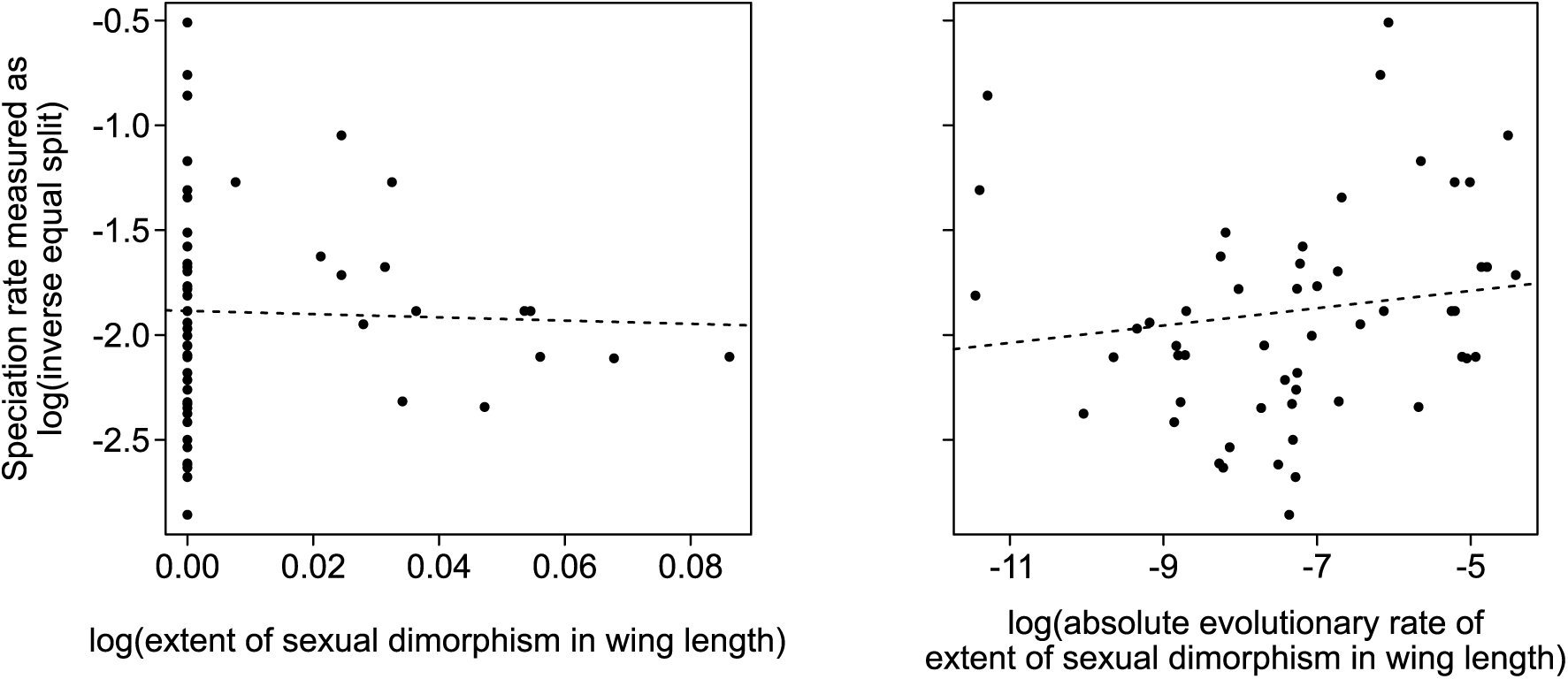
Speciation rate in relation to the extent of sexual dimorphism in wing length, measured here as log((male wing length)/(female wing length)), as depicted in the left panel and in relation to the evolutionary change of the extent of sexual dimorphism in wing length, estimated by absolute evolutionary rates in log((male wing length)/(female wing length)) using the R function “RRphylo” in the package “RRphylo” after log-transformation (see text), as depicted in the right panel. Here, estimates using one of 1000 phylogenetic trees with simple regression lines were depicted for illustrative purpose (broken lines were used for non-significant correlations; see text for formal analyses using 1000 trees)

